# Functional validation of a finding from a mouse genome-wide association study demonstrates that a mutant allele of *Azi2* alters sensitivity to methamphetamine

**DOI:** 10.1101/2020.08.22.262709

**Authors:** Xinzhu Zhou, Amanda Barkley-Levenson, Patricia Montilla-Perez, Francesca Telese, Abraham A. Palmer

**Author notes:** **Corresponding Author:** Biomedical Research Facility II (BRF2), Room 3A24; 9500 Gilman Dr.; Mail Code 0667; La Jolla, CA 92093-0667.

## Abstract

Methamphetamine is a widely abused psychostimulant. In a previous genome-wide association study (**GWAS**), we identified a locus that influenced the stimulant response to methamphetamine. That locus was also an eQTL for the gene *Azi2*. Based on those findings, we hypothesized that heritable differences in the expression of *Azi2* were causally related to the differential response to methamphetamine. In this study, we created a mutant *Azi2* allele that caused lower *Azi2* expression and enhanced the locomotor response to methamphetamine; however, based on the GWAS findings, we had expected lower *Azi2* to decrease rather than increase the stimulant response to methamphetamine. We then sought to explore the mechanism by which *Azi2* influenced methamphetamine sensitivity. A recent publication had reported that the 3’UTR of *Azi2* mRNA downregulates the expression of *Slc6a3*, which encodes the dopamine transporter (**DAT**), which is a key target of methamphetamine. We evaluated the relationship between *Azi2*/*Azi2* 3’UTR and *Slc6a3* expression in the VTA in the mutant *Azi2* mice and in a new cohort of CFW mice. We did not observe any correlation between *Azi2* and *Slc6a3* in the VTA in either cohort. However, RNA sequencing confirmed that the *Azi2* mutation altered *Azi2* expression and also revealed a number of potentially important genes and pathways that were regulated by *Azi2*, including the metabotropic glutamate receptor group III pathway and nicotinic acetylcholine receptor signaling pathway. Our results support a role for *Azi2* in methamphetamine sensitivity; however, the exact mechanism does not appear to involve regulation of *Slc6a3* and thus remains unknown.

## Introduction

Methamphetamine is a widely abused psychomotor stimulant. In the United States, approximately 1.6 million people reported using methamphetamine in the past year (1). Methamphetamine can produce feelings of euphoria, heightened energy, and enhanced focus. Although most users experience stimulants such as methamphetamine, amphetamine and cocaine as pleasurable, studies in both humans and animals have found marked individual differences (2–7) which may be at least partially genetic (8,9). These differences are believed to alter the behavioral and subjective response to methamphetamine and may therefore alter risk for methamphetamine abuse. Such individual differences are often modeled by studying the acute locomotor stimulant response to drugs in rodents (10,11)

Over the last decade, large scale genome-wide association studies (**GWAS**) have facilitated the identification of thousands of loci that influence complex traits including abuse of alcohol and other substances (Buchwald et al., 2020; Kinreich et al., 2019; Liu et al., 2019; Sanchez-Roige et al., 2019; Walters et al., 2018; Zhou et al., 2020). GWAS in model organisms provide a complementary approach to human GWAS and have also identified loci for numerous traits, including several that are related to drug abuse (*e.g*. (19,20)). An advantage of GWAS in model organisms is that putatively identified genes can be directly manipulated to assess causality and to better understand the underlying molecular mechanisms; however, such manipulations are not frequently performed.

We previously reported a GWAS that examined a number of behavioral and physiological traits, including the acute locomotor stimulant response to methamphetamine using 1,200 outbred Carworth Farms White (**CFW**) male mice (20). CFW mice are a commercially available outbred population that have relatively small linkage disequilibrium blocks, which allow the identification of small loci and thus narrow the number of genes that might be causally related to the trait being measured (20). One of the many genome-wide significant findings from that study was an association between a locus on chromosome 9 (rs46497021) and the locomotor response to methamphetamine injection. That locus also contained a *cis*-expression quantitative trait locus (***cis*-eQTL**) for the gene 5-azacytidine–induced gene 2 (***Azi2***) in the striatum (the peak eQTL SNP was rs234453358, which is in strong LD with rs46497021). Based on these data, we suggested that heritable differences in *Azi2* expression may be the molecular mechanism by which that locus influenced the acute locomotor response to methamphetamine. However, in our prior publication we did not directly test that hypothesis by manipulating *Azi2* expression.

*Azi2* is known to activate NFκB (21), to be involved in TNF-induced cell death (22–25), and to influence immune response (26). However, its role in methamphetamine sensitivity remains unclear. One possible mechanism by which *Azi2* could influence the response to methamphetamine was proposed by another group after the publication of Parker et al (2016) (20). Liu et al (2018) identified the second half of 3’ UTR of *Azi2* mRNA (^**AZI2**^**3’UTR**) as a regulator of *Slc6a3* (Liu et al., 2018), which encodes the dopamine transporter (**DAT**), a critical regulator of the neurotransmitter dopamine concentrations in the synaptic cleft. Importantly, DAT is the molecular target of methamphetamine (28), providing a plausible mechanism by which *Azi2* might influence sensitivity to methamphetamine. In addition to a number of cellular assays, Liu et al (2018) also showed that rats that had been bi-directionally selected for alcohol preference showed differential ^AZI2^3’UTR expression and differential expression of *Slc6a3* expression in the ventral tegmental area (**VTA**) (Liu et al., 2018).

To test the hypothesis that *Azi2* was the gene responsible for the association detected in our GWAS, and that its action was mediated via *Slc6a3*, we created a mutant *Azi2* allele using CRISPR/Cas9 to generate a frameshifting mutation in exon 3 of *Azi2*. Using the *Azi2* KO mice, we examined the effect of this mutant allele on the acute locomotor response to methamphetamine. We evaluated gene expression in the striatum, the brain tissue in which the eQTL for *Azi2* expression was identified in Parker et al (2016) (20), to validate the elimination of *Azi2*. In an effort to determine whether our mutant allele altered the expression of *Azi2* 3’UTR, and whether such changes might alter *Slc6a3*, as predicted by Liu et al (2018), we also measured gene expression in the VTA, since DAT is expressed there. We also performed parallel studies in a new cohort of CFW mice to confirm that the allele identified in Parker et al (2016) was indeed associated with changes in *Azi2* and to determine whether it also associated with differential *Slc6a3* expression in the VTA, a brain region that was not examined in Parker et al (2016).

## Materials and Methods

### Establishment of an *Azi2* knockout mouse line using CRISPR/Cas9

We followed the JAX protocol of microinjection of CRISPR mix using sgRNA and *Cas9* mRNA (https://www.jax.org/news-and-insights/1998/july/superovulation-technique). We designed a sgRNA targeting exon 3 of *Azi2* (Vector Name: pRP[CRISPR]-hCas9-U6>{20nt_GGGCCGAGAACAAGTGAATA}; Table S1).

All animal procedures were approved by the local Institutional Animal Care and Use Committee and were conducted in accordance with the NIH Guide for the Care and Use of Laboratory Animals. The CRISPR microinjection procedures were performed at the University of California San Digeo, Moores Cancer Center, Transgenic Mouse Core. We ordered five C57BL/6J stud males (7-8 weeks old) and five C57BL/6J females (3-4 weeks old) from the Jackson Laboratory (Bar Harbor, ME). Upon arrival at the vivarium, the stud males were singly-housed and the females were housed in groups of four. On Day 1 of the microinjection week, all five females were super-ovulated via 0.1ml pregnant mare serum (PMS) intraperitoneal injection per animal. On Day 3, all females were super-ovulated via 0.1ml human chorionic gonadotropin (HCG) intraperitoneal injection per animal. After hormonal priming, each female was placed into the home cage of one stud male for mating. On Day 4, ovulation was expected to occur, and females were separated from the stud males. Fallopian tubes were dissected out from the mated females and were collected in M2 medium. Zygotes were harvested and injected with the CRISPR mix (625ng = 3.1ul×200ng/ul of *Azi2* sgRNA + 1250ng = 5ul×250ng/ul of *Cas9* mRNA + 16.9ul ph7.5 IDTE; total volume 25ul). Injected zygotes were surgically transplanted to pseudopregnant female C57BL/6JOlaHsd (Harlan) mice. Pregnant surrogate dams were singly caged one week before the expected birth date of the pups. C-sections were carried out if necessary.

### *Azi2* knockout (KO) line breeding and genotyping scheme

We obtained 14 *Azi2* CRISPR founders (five males, nine females). The founders were genotyped via Sanger sequencing to verify the presence of deletions. The male founders were then backcrossed to wildtype C57BL/6J mice to minimize the effect of off-targeting. F_1_/F_2_ *Azi2* KO mice were genotyped via Sanger/NGS to ensure the transmission of the mutant allele. Heterozygous F_1_s were paired to produce F_2_s, which were genotyped via next-generation sequencing. Among others, we identified one 7bp deletion in F_2_s. This deletion was predicted to cause mRNA degradation of the four full-length RefSeq supported *Azi2* transcripts, ENSMUST00000044454.11, ENSMUST00000133580.7, ENSMUST00000134433.7, and ENSMUST00000154583.7, and three shorter, predicted *Azi2* transcripts, ENSMUST000000143024.1, ENSMUST000000130735.7, and ENSMUST000000127189.1. We genotyped this deletion via restriction fragment length polymorphism (**RFLP**); this mutation harbors the *StuI* restriction enzyme target site, which allows us to easily genotype the mice. The sgRNA for the CRISPR/Cas9 procedure and the genotyping primers for *Azi2* are included in Table S1.

In this study, we will refer to our version of the *Azi2* mutation as the ‘mutant *Azi2* allele’, all progeny of the *Azi2* KO founders as the ‘Azi2 *KO* mice’, and the homozygous mutant *Azi2* mice as the ‘mutant *Azi2* mice’.

### Locomotor response to methamphetamine

We assessed the locomotor response to methamphetamine using the protocol previously described in Parker et al 2016 (20). Adult male and female mice were tested over a three-day period between 0800 and 1700 h. Mice were group housed 2-5 per cage on a 12h/12h light-dark cycle with lights on at 0600 h. Mice were transported to the procedure room at least 30 min before testing, which allowed them to habituate to the new environment while remaining in their home cages. On each day of testing, each animal was briefly placed in an individual clean cage. Animals were weighed to determine the volume of injection (0.01 ml/g body weight). On day one and day two, mice received an i.p. injection of 0.9% saline solution; on day three, mice received an i.p. injection of methamphetamine solution (1.5 mg/kg of (+)-Methamphetamine hydrochloride; Sigma Life Science, St. Louis, MO, dissolved in a 0.9% saline solution). Immediately following injection, each mouse was placed in the test chamber for activity recording. All animals were measured using the Versamax software (AccuScan Instruments, Columbus, OH). At the end of the 30 min test, mice were returned to their home cages. Test chambers were sprayed with 10% isopropanol between tests. At the end of each test day, animals were returned to the vivarium.

We removed one heterozygote and one mutant mouse whose locomotor activity on day 3 were more than three standard deviations away from the mean. We analyzed a total of 135 mice and the ratio of genotypes was consistent with expectations: 33 wildtype littermates (18 females, 15 males), 67 heterozygotes (31 females, 36 males), and 35 mutants (18 females, 17 males).

### Genotyping CFW naïve mice

CFW mice were genotyped via Sanger sequencing. Genotyping primers for CFW mice are included in Table S5. We genotyped the GWAS top SNP for the trait “Distance traveled, 0–30 min, on day 3” at rs46497021 (rs46497021) and the eQTL top SNP for *Azi2* expression in the striatum at rs234453358 (rs234453358; see Parker et al (2016) (20)).

### Brain tissue collection

Mouse brain tissue was dissected using an adult mouse brain slicer matrix with 1.0 mm coronal section slice intervals (ZIVIC instruments, Pittsburgh, PA, USA). Striatum was collected from slice Bregma 0 to 2 and VTA was collected from slice Bregma −4 to −2. Four tissue punches, two on the left and two on the right hemisphere, were collected for each animal. After dissection, brain tissue was placed in an Eppendorf tube that was fully submerged in dry ice.

### Analysis of CFW data from Parker et al (2016)

Because genotypes in Parker et al (2016) were represented as genotype probabilities, we first converted probabilities to dosages and then coded dosages < 0.2 as homozygous reference, dosages > 0.8 and <1.2 as heterozygous, dosages > 1.8 homozygous alternative. A hundred and seven mice with intermediate dosage values are excluded from the plots. We used likelihood ratio test of nested models (*lmtest* R package; (29)) to examine the genotype effect.

### qRT-PCR

Primers and probes selected for *Azi2*, 3’UTR of *Azi2*, *Slc6a3*, and *Gapdh* gene expression assays are shown in Table S2. We used TaqMan gene expression assays for *Azi2*, *Slc6a3*, and *Gapdh*. We custom-designed the gene expression assay for the 3’UTR of *Azi2*. We custom designed the TaqMan primers and the FAM-MGB probe for 3’UTR of *Azi2* according to the Custom TaqMan Assay Design Tool (https://tools.thermofisher.com/content/sfs/manuals/cms_042307.pdf). We ran all qRT-PCR experiments in duplicate on the StepOnePlus™ Real-Time PCR System (Applied Biosystems, Waltham, MA, USA).

To demonstrate that the 7bp deletion led to the degradation of full-length *Azi2* mRNA transcripts, we performed qRT-PCR that amplified the exon 6-7 junction in *Azi2* mRNA transcripts ENSMUST00000044454.11, ENSMUST00000133580.7, and ENSMUST00000134433.7; in ENSMUST00000154583.7 this same sequence corresponds to exon 5-6. This amplicon would detect the four RefSeq *Azi2* transcripts and three predicted transcripts ENSMUST00000135251.1, ENSMUST00000130735.7, and ENSMUST00000133814.1. Our CRISPR/Cas9 deletion scheme ensured that all four full-length, RefSeq supported *Azi2* transcripts and a few short *Azi2* predicted transcripts would be degraded via nonsense-mediated mRNA decay due to the deletion. Given the mRNA degradation, our qRT-PCR design for whole-gene *Azi2* expression would only detect the two short, predicted *Azi2* transcripts, ENSMUST0000013525.1 and ENSMUST00000133814.1, in heterozygous and mutant mice.

The *Azi2* 3’UTR on exon 8 that we amplified using qRT-PCR is homologous to the 3’UTR amplified in the rat alcohol model (Liu et al., 2018) and is only present in two of the four full-length *Azi2* transcripts, ENSMUST0000044454.11 and ENSMUST00000133580.7.

Using qRT-PCR, we measured *Azi2* expression in 44 mice from the *Azi2* KO line, which included 15 homozygous mutants. We measured the *Azi2* 3’UTR expression in an additional 33 mice from the *Azi2* KO line. Each batch of the *Azi2* KO mice used for gene expression assays were of similar age (199-201 days at sacrifice for the mice used for measuring *Azi2* mRNA and 226-232 days at sacrifice for the mice used for measuring *Azi2* 3’UTR mRNA).

We also performed qRT-PCR in CFW mice. We removed three animals whose *Azi2* or *Slc6a3* gene expression was more than three standard deviations away from the mean. We analyzed a final set of 31 CFW mice for the *Azi2* and *Slc6a3* expression in the striatum and in the VTA (rs46497021: ‘GG’ n=5, ‘GA’ n=20, ‘AA’ n=6; rs234453358: ‘AA’ n=12, ‘AG’ n=13, ‘GG’ n=6).

### RNA-Sequencing

We extracted RNA from the striatum and VTA of the mouse brain and prepared cDNA libraries from 68 samples with RNA integrity scores ≥7.0 (32 *Azi2* KO mice: 9 wildtype, 13 heterozygous, 10 mutant; 36 CFW mice: 12 ‘GG’, 12 ‘GA’, 12 ‘AA’ at rs46497021, 12 ‘AA’, 12 ‘AG’, 12 ‘GG’ at rs234453358) as measured on a TapeStation (Agilent, Santa Clara, CA, USA). The cDNA libraries were prepared with the NEBNext^®^ Ultra™ II Directional RNA Library Prep Kit for Illumina (NEW ENGLAND BioLabs, Ipswich, MA, USA) and sequenced on two lanes (two chips, one lane on each chip) of an Illumina NovaSeq S4 using 100◻bp, paired-end reads.

All sequencing reads passed the Illumina sequencing quality score of 20. We used HISAT2 (30) to align the adapter-trimmed paired-end reads simultaneously to mouse reference genome mm10. We used HTSeq to assign reads to gene features, in which the union of all the sets of all features overlapping each position *i* in the read was counted (31). We then examined potential expression outliers due to technical variance in PCA plots and removed a wildtype VTA sample from the *Azi2* KO cohort, a ‘GG’ VTA sample at rs234453358 and an ‘AA’ striatum sample at rs234453358 from the CFW cohort. We used the final set of 31 *Azi2* KO samples (16 striatum samples: 5 wildtype, 6 heterozygous, 5 mutant; 15 VTA samples: 3 wildtype, 7 heterozygous, 5 mutant) and 34 CFW samples (17 striatum samples: 5 ‘AA’, 6 ‘AG’, 6 ‘GG’ at rs234453358; 17 VTA samples: 6 ‘AA’, 6 ‘AG’, 5 ‘GG’ at rs234453358).

We calculated the read counts aligned to each exon feature of *Azi2* by providing Samtools the genomic coordinates of the exon features (32). Then, we normalized the read counts by dividing the raw reads by the length of the exon feature and the total number of reads in the sample. Normalized read counts for *Azi2* were calculated by summing normalized read counts aligned to exons 1-8 for transcripts ENSMUST00000044454.11, ENSMUST00000133580.7, and ENSMUST00000134433.7 and exons 1-7 for transcript ENSMUST00000154583.7 (chr9.118040522-chr9.118069794). Normalized read counts for *Azi2* 3’UTR were calculated using the region from chr9.118063214-chr9.118063336, and mouse genomic region that is homologous to the second half of *Azi2* 3’UTR in rats, which matches the procedure described in Liu et al (2018) (27).

We used DESeq2 to perform differential expression analysis (33). Prior to count normalization and differential expression analysis, we calculated the average count per million (CPM) within each cohort and tissue combination across genes and samples. We only retained genes with CPM larger than 1. Then, we disabled independent filtering, which identifies the maximum number of adjusted *p* values lower than a significance level alpha based on the mean of normalized counts. We kept the “Cook’s distance” parameter in DESeq2, which removes genes with extreme count outliers that do not fit well to the negative binomial distribution. These procedures ensured that genes with extremely low and large raw counts are removed and the same set of genes are used in differential expression analysis in between-genotype comparisons. At the end of filtering steps, we had 14,680 genes in *Azi2* KO striatum, 15,003 genes in *Azi2* KO VTA, 14,594 genes in CFW striatum, 14,936 genes in CFW VTA. All differential expression analyses performed on the *Azi2* KO mice used the design ~ genotype + sex because we observed a separation by sex effect in PCA plots; analyses on the CFW mice only included the genotype factor because all mice were male.

## Results

### Creation and characterization of mutant *Azi2* allele using CRISPR/Cas9

To investigate whether *Azi2* might be the gene underlying the association between the locus on chromosome 9 and the locomotor stimulant response to methamphetamine that we had previously observed in CFW mice (20), we created a mutant allele of *Azi2*. Because of technical difficulties generating embryos from CFW, and because of the more complicated breeding programs necessary for maintaining a mutant allele on an outbred background, we generated the mutant alleles on the C57BL/6J background. We designed a sgRNA targeting exon 3 of *Azi2* (Figure 1a; Table S1), because exon 3 harbors the start codon of *Azi2* and is included in all four annotated transcripts of *Azi2*. We selected a mutant mouse line that carried a 7bp frameshifting deletion in exon 3 of *Azi2* (Table S2).

**Figure 1.**
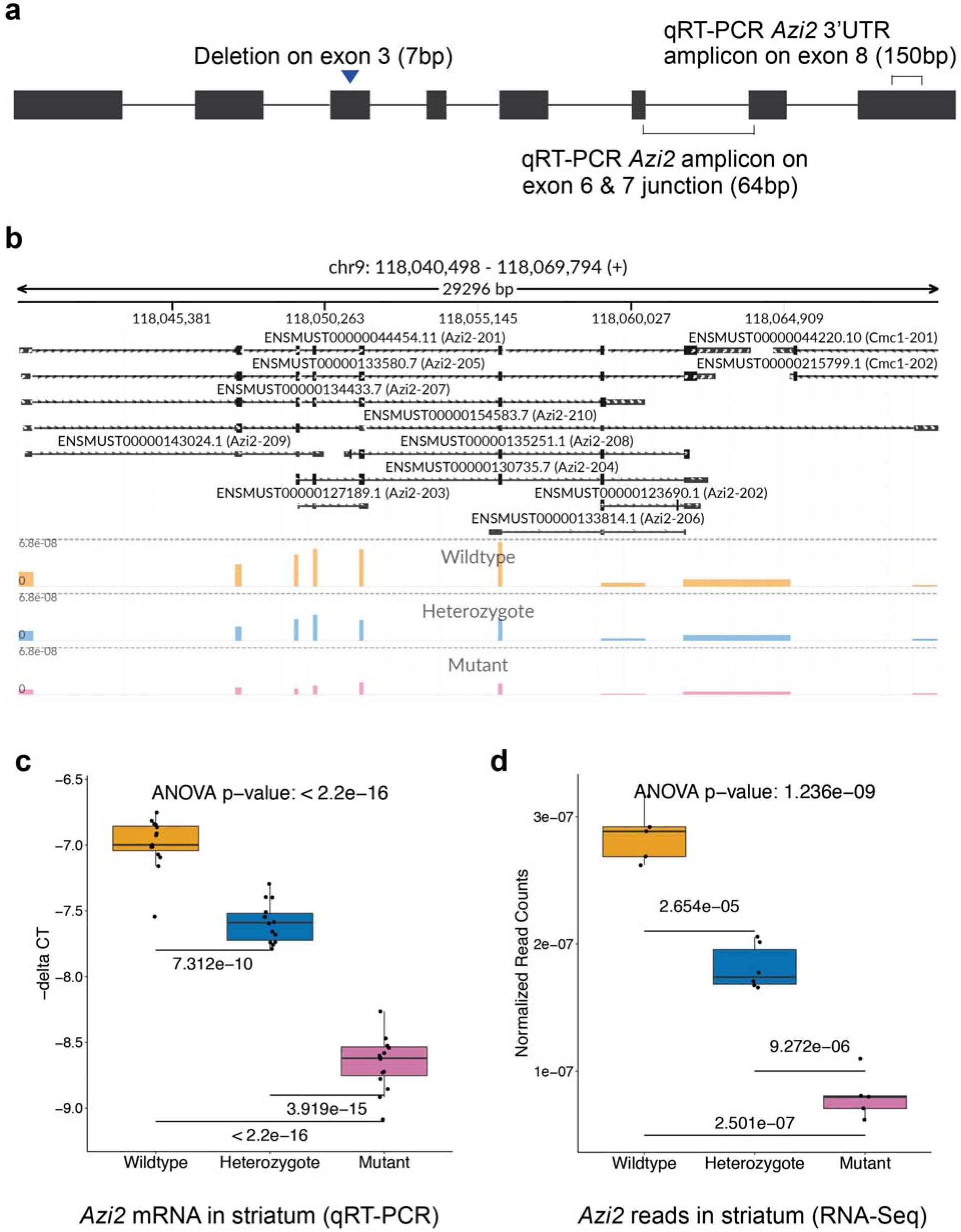
A 7bp deletion on exon 3 of *Azi2* was generated by CRISPR/Cas9. The genomic position of the CRISPR/cas9 deletion and the qRT-PCR amplicons for *Azi2* mRNA and *Azi2* 3’UTR mRNA are indicated in **a**. **b** RNA-Seq reads aligned to the exons of *Azi2* in a wildtype, a heterozygous, and a mutant mouse in the *Azi2* KO line show the effect of the CRISPR/Cas9 deletion on mRNA abundance in the striatum. The mutant mouse had fewer reads across all the exons than the heterozygote, which in turn had fewer reads than the wildtype. The RNA-Seq reads aligned to each exon feature were normalized to the length of exon and the total read counts of the sample. To choose the most representative sample for each genotype for *Azi2* expression in striatum, we calculated the average normalized read counts for each genotype and identified the sample closest to the average. Note that the tracks include all *Azi2* transcripts annotated in the comprehensive gene annotation file of GRCm38.p6; only the top four transcripts are supported by RefSeq. The genome tracks were plotted using the Python visualization tool svist4get (37). **c** Using qRT-PCR, we showed that *Azi2* expression in the striatum in mutant *Azi2* KO mice is significantly lower than the heterozygous and the wildtype mice (*F*(2,41) = 319.41, p < 2.2×10^−16^). Delta CT was calculated as the mean CT of target gene (*Azi2*) – the mean CT of the control gene (*Gapdh*); larger values of – delta CT indicate higher gene expression level. *Azi2* expression was measured using real time PCR in 15 wildtype, 14 heterozygous, and 15 mutant *Azi2* KO mice. We used Welch two sample t-test to make between group comparisons, which show that all groups were different (wildtype vs heterozygote *t*(26.301) = −9.365, p = 7.312×10^−10^; heterozygote vs mutant *t*(25.906) = −16.297, p = 3.919×10^−15^; wildtype vs mutant *t*(27.944) = −23.3, p < 2.2×10^−16^). **d** Using RNA-Seq we showed that normalized read counts mapped to *Azi2* in the striatum of the *Azi2* KO line also show a significant effect of the CRISPR/Cas9 deletion (*F*(2,13) = 146.04, p = 1.236×10^−9^). RNA-Seq was performed in 5 wildtype, 6 heterozygous, and 5 mutant *Azi2* KO striatum samples. Between group comparisons show that all groups are different (wildtype vs heterozygote *t*(7.8331) = 8.7235, p = 2.654×10^−5^; heterozygote vs mutant *t*(8.537) = 9.3156, p = 9.272×10^−6^; wildtype vs mutant *t*(7.8068) = 16.399, p = 2.501×10^−7^).

Because Parker et al (2016) had identified a coincident eQTL for *Azi2* in the striatum, we sought to confirm that the mutant allele would reduce *Azi2* expression in that same brain region. In addition, because the Liu et al study (Liu et al., 2018) study focused on the role of *Azi2* in the VTA, we also examined *Azi2* expression in that brain region. Using qRT-PCR in wildtype, heterozygote and mutant *Azi2* KO mice, we confirmed that the 7bp deletion led to significantly decreased abundance of *Azi2* mRNA in the striatum (Figure 1c, Figure S1). In a separate cohort of mice, we used RNA-Seq in the *Azi2* KO mice to further demonstrate that that the 7bp deletion led to significantly decreased abundance of *Azi2* mRNA in the striatum (Figure 1b&d, Figure S2a). In addition to the decreased mRNA abundance associated with the mutant allele, many of the remaining transcripts will be frameshifted and thus will not encode functional protein because they.

### The locomotor stimulant response to methamphetamine was greater in mutant *Azi2* mice

Having created a mutant *Azi2* allele, we next sought to examine whether mutant *Azi2* mice showed an altered locomotor response to methamphetamine. In particular, we sought to precisely replicate the protocol used in Parker et al (2016), in which saline was given on days 1 and 2 and 1.5 mg/kg methamphetamine was given on day 3. Parker et al. (2016) found that mice with more ‘A’ alleles at rs46497021 exhibited greater sensitivity to methamphetamine; data from that paper are replotted in Figure 2a-c. We also reanalyzed data from Parker et al. (2016) to demonstrate that *Azi2* expression was higher in individuals that had the ‘A’ allele at rs46497021 (Figure S3a). The same was also true at rs234453358, which was in LD with rs46497021 (Figure S3b). In our *Azi2* KO line, we found that locomotor responses to saline on days 1 and 2 did not differ among the three genotype groups (Figure 2d, e), but the response to methamphetamine (day 3) was different, with the mutant mice (lower *Azi2* expression) showing higher methamphetamine sensitivity (Figure 2f; Figure S4). Surprisingly this direction of effect is opposite to that of the CFW mice from Parker et al (2016) where mice with more ‘A’ alleles at rs46497021 had higher *Azi2* expression and higher methamphetamine sensitivity (Figure 2c).

**Figure 2a-f.**
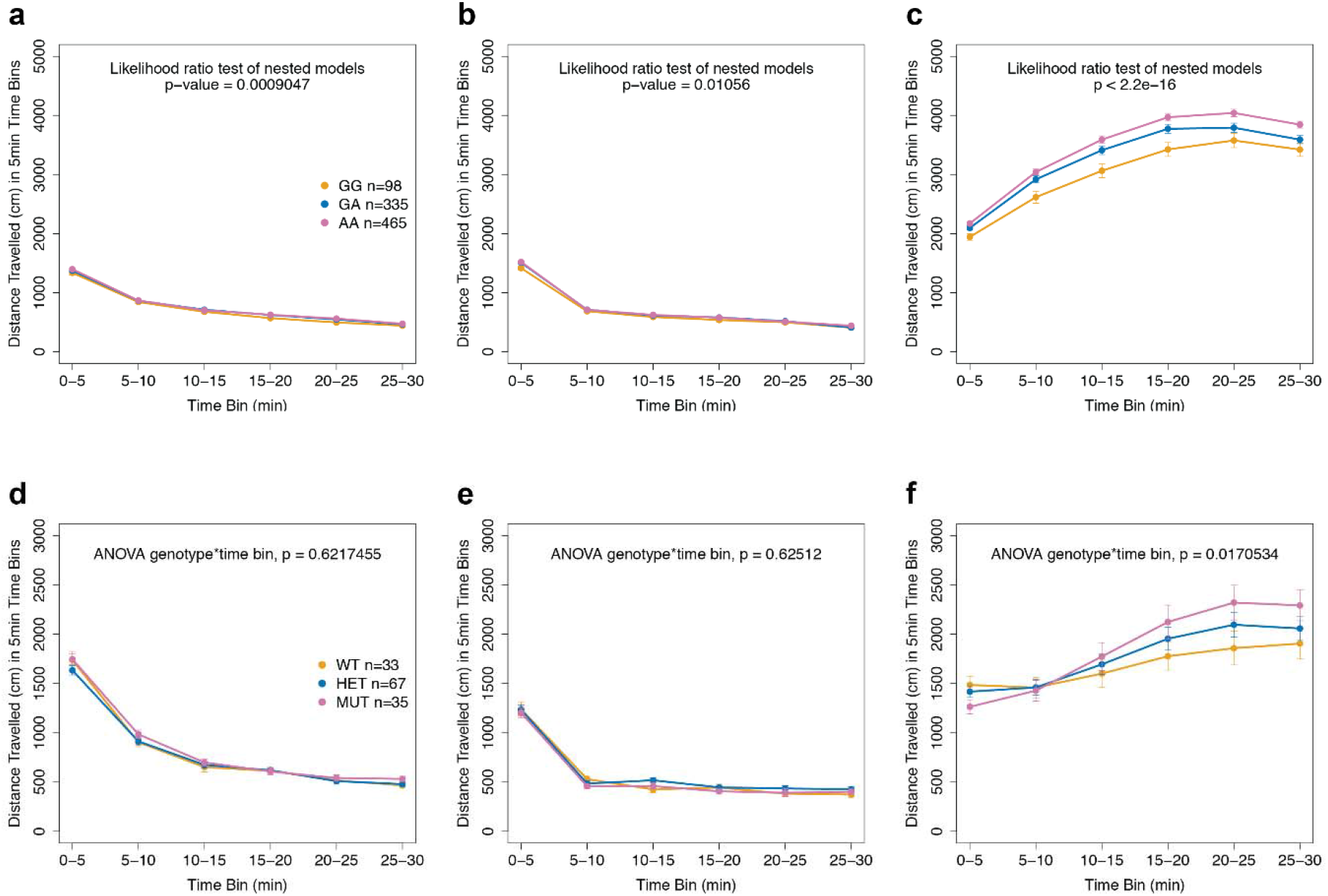
Locomotor response to methamphetamine was moderately heightened in mutant *Azi2* KO mice. **a – c** Locomotor activity of CFW (n=898 as previously reported in Parker et al 2016) following administration of saline (days 1 and 2) and methamphetamine (day 3), plotted in 5 min time bins. The genotype effect was noticeably more significant on day 3 of methamphetamine injection (*X*^2^(1,2) = 70.598, p < 2.2×10^−16^) than that on day 1 (*X*^2^(1,2) = 11.013, p = 0.0009047) and day 2 of saline injection (*X*^2^(1,2) = 6.5381, p = 0.01056). **d** – **f** A total of 135 mice from the *Azi2* KO line were tested in the locomotor response to methamphetamine experiment. To evaluate the effect of the mutant *Azi2* allele, we used an ANOVA to analyze the effects of genotype, time bin and sex on the locomotor response to methamphetamine. The 3-way interaction was not significant (see Table S3); however, there was a significant interaction between genotype and time bin (*F*(2,798)= 4.09230, p = 0.0170534). Post-hoc tests did not identify any particular time bin that was different, though there were some trends towards differences between the wildtype and mutant mice (Tables S3 and S4).

### *Azi2* mRNA and *Azi2* 3’UTR mRNA did not downregulate *Slc6a3* expression in *Azi2* KO mice

Liu et al (2018) reported that the 3’UTR of *Azi2* mRNA negatively regulates *Slc6a3* in the midbrain of rats, and that there was an increase in expression of the 3’UTR of *Azi2* in the non-alcohol preferring rats compared to alcohol preferring rats (Liu et al., 2018). Based on those data, Liu et al (2018) argued that regulation of *Slc6a3* expression by the 3’UTR of *Azi2* is important for substance use related traits. Based on these data, we hypothesized that *Azi2* expression in CFW mice might have led to altered response to methamphetamine because of its ability to regulate *Slc6a3*.

We tested this hypothesis using both qRT-PCR and RNA-Seq. Using qRT-PCR, we measured the level of *Azi2*, *Azi2* 3’UTR and *Slc6a3* mRNA in the VTA (Figure 1a). The *Azi2* 3’UTR on exon 8 that we amplified using qRT-PCR is homologous to the 3’UTR amplified in the rat alcohol model (Liu et al., 2018) and is only present in two of the four full-length *Azi2* transcripts, ENSMUST0000044454.11 and ENSMUST00000133580.7 (Figure 1b). We found that the expression of both *Azi2* and *Azi2* 3’UTR amplicons were decreased in a genotype-dependent manner in the VTA in the mutant mice (Figure S1a; Figure S5a). However, there was no significant effect of genotype on *Slc6a3* expression in the VTA (Figure S1b; Figure S5b). Furthermore, we did not observe any correlation between the expression of *Azi2*, *Azi2* 3’UTR or *Slc6a3* in the VTA (Figure 3a&b).

**Figure 3a-f.**
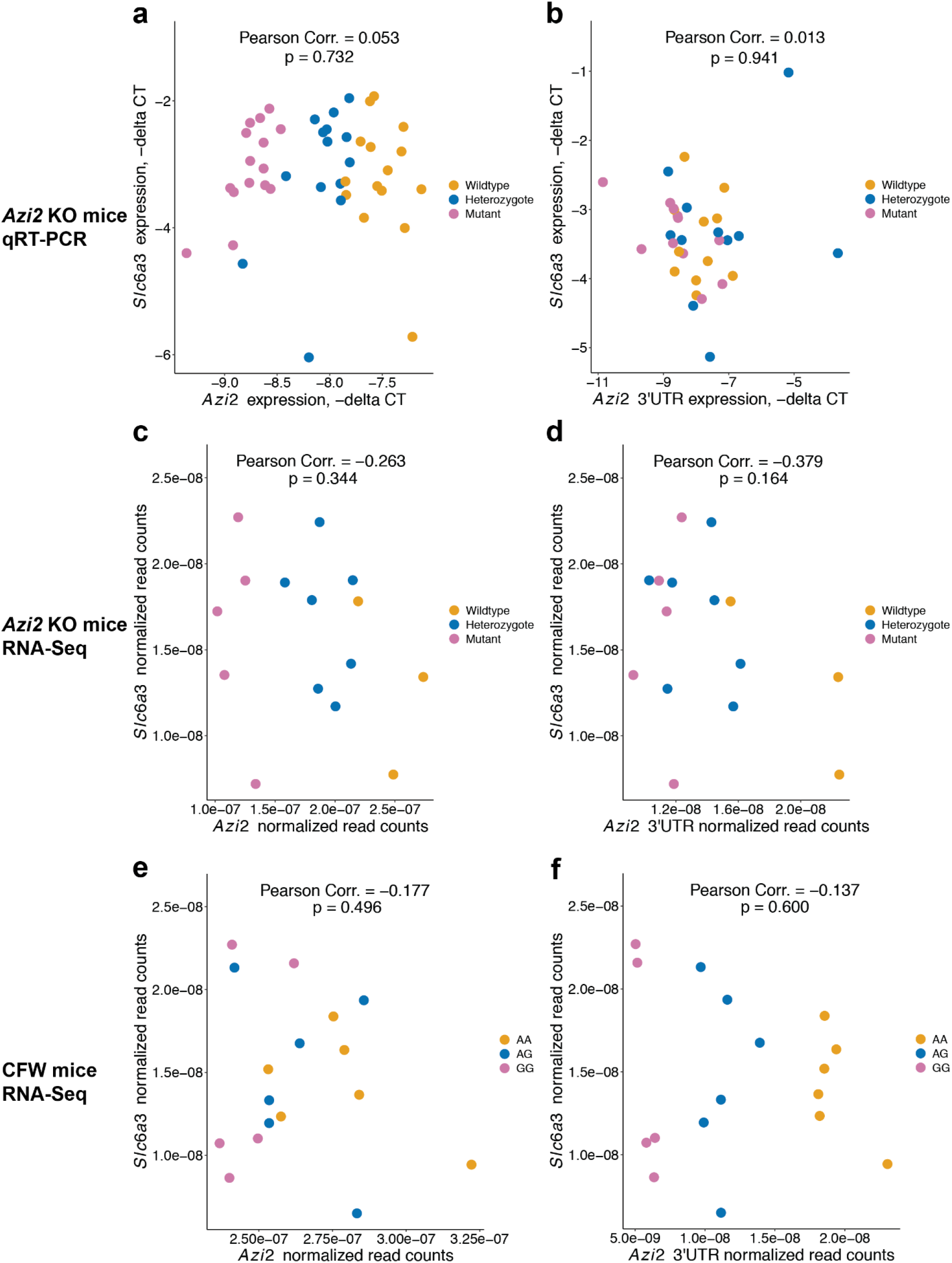
*Azi2* and *Azi2* 3’UTR did not down-regulate *Slc6a3* in the VTA in the *Azi2* KO mice; *Azi2* did not down-regulate *Slc6a3* in the VTA of naïve CFW mice. We used **a** 44 *Azi2* KO mice (wildtype = 15, heterozygote= 14, mutant= 15) for examining the correlation between *Azi2* and *Slc6a3* expression measured by qRT-PCR, **b** 33 *Azi2* KO mice (wildtype = 11, heterozygote= 11, mutant= 11) for examining the correlation between *Azi2* 3’UTR and *Slc6a3* expression measured by qRT-PCR, **c** & **d** 15 *Azi2* KO mice (wildtype = 3, heterozygote= 7, mutant= 5) for examining the correlation between *Azi2*/*Azi2* 3’UTR and *Slc6a3* normalized read counts measured by RNA-Seq, and **e** & **f** 17 CFW mice (‘AA’ = 6, ‘AG’ = 6, ‘GG’ = 5) for examining the correlation between *Azi2*/*Azi2* 3’UTR and *Slc6a3* normalized read counts measured by RNA-Seq. Neither *Azi2* nor *Azi2* 3’UTR expression was negatively correlated to *Slc6a3* expression in the VTA in either cohort of mice. **a & b** Neither the level of *Azi2* (*r*(42) = 0.05321076, p = 0.7316) nor *Azi2* 3’UTR expression (*r*(31) = 0.01339348, p = 0.941) was negatively correlated to the expression of *Slc6a3* in the VTA. **c & d** No significant correlation between *Azi2* and *Slc6a3* (*r*(13) = −0.2627568, p = 0.3441) or *Azi2* 3’UTR and *Slc6a3* (*r*(13) = −0.3786666, p = 0.164) was observed. **e & f** Neither *Azi2* (*r*(15) = −0.1771307, p = 0.4964) nor *Azi2* 3’UTR (*r*(15) = −0.1370425, p = 0.5999) was negatively correlated with *Slc6a3* expression in the VTA at the eQTL top SNP.

We also used RNA-Seq to examine the hypothesis that *Azi2*/*Azi2* 3’UTR could downregulate *Slc6a3* in the VTA in the *Azi2* KO line. Expression of *Azi2* and *Azi2* 3’UTR were lower in the heterozygote and mutant mice; however, there was no effect of genotype on *Slc6a3* expression (Figure 3c&d). Taken together, our results do not support the negative correlation between *Azi2*/*Azi2* 3’UTR and *Slc6a3* in the VTA in our *Azi2* KO mice.

### *Azi2* mRNA did not downregulate *Slc6a3* regulation in naïve CFW mice in VTA

We also examined the relationship between *Azi2*, *Azi2* 3’UTR and *Slc6a3* in a behaviorally and drug naïve CFW mice to address the possibility that the strain difference between C57BL/6J and CFW may have contributed to the previously reported negative correlation between *Azi2*/*Azi2* 3’UTR and *Slc6a3* expression. Using 31 male CFW mice, we found that there was no correlation between *Azi2* nor *Azi2* 3’UTR and *Slc6a3* in the VTA (Figure 3e&f; Figure S6; Figure S7).

### Analysis of gene expression differences using RNA-Seq in *Azi2* KO and in CFW mice

Next, we sought to identity genome-wide changes in gene expression observed in the *Azi2* KO line using the RNA-Seq data. When comparing wildtype to mutant mice, we identified 23 differentially expressed genes in the striatum at FDR < 0.1 (Figure 4a; Table S6). In the VTA, the same comparison identified four differentially expressed genes (Figure 4b; Table S6). For both tissues, *Azi2* was by far the most significantly differentially expressed gene. When comparing wildtype to heterozygous mice, we found that *Azi2* was the only differentially expressed gene in both the striatum and the VTA (Figure S8a, S8b).

**Figure 4a-d.**
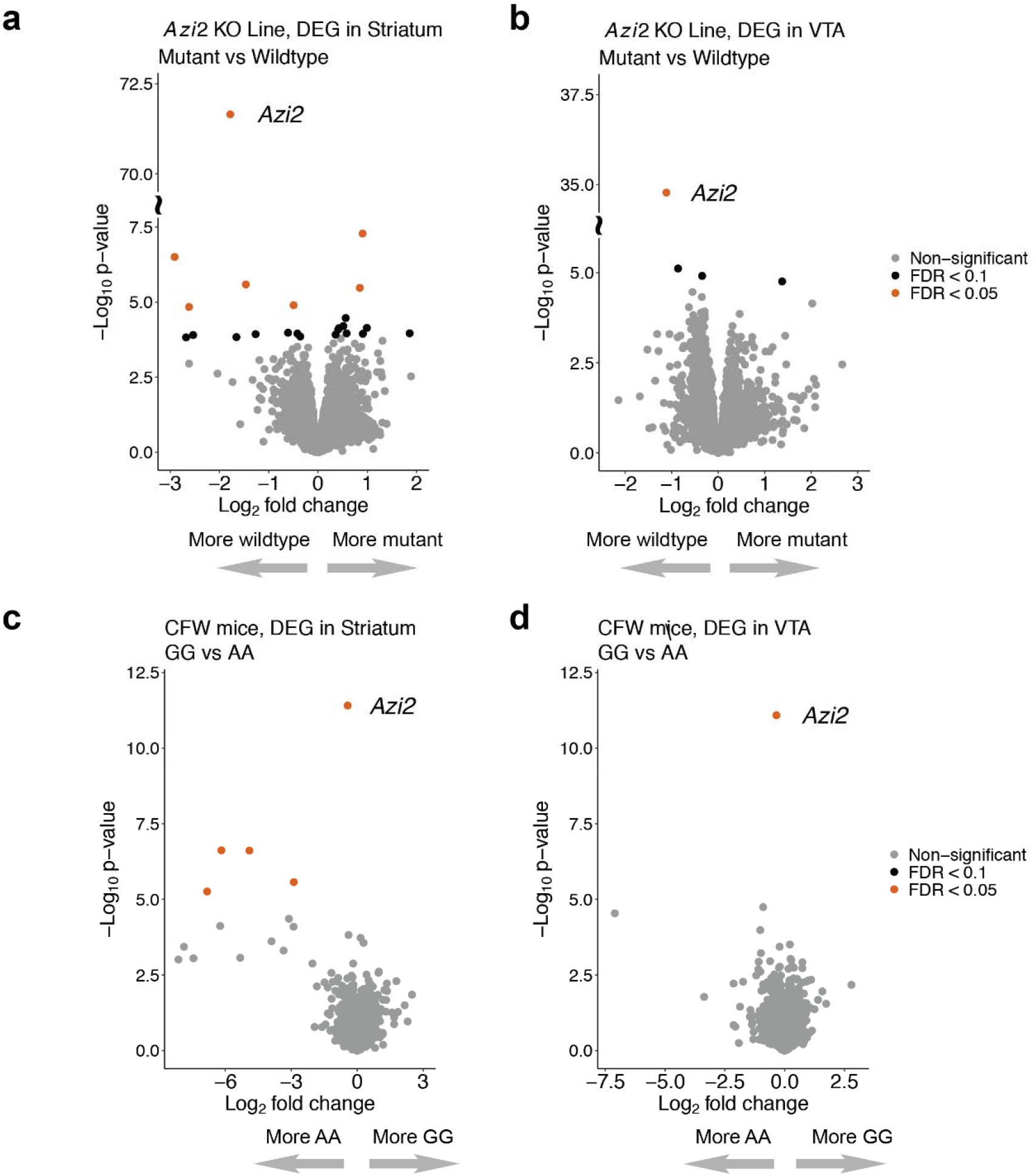
*Azi2* (ENSMUSG00000039285.12) was consistently differentially expressed between mutant vs wildtype mice in the *Azi2* KO line and between homozygous alternative (‘GG’) vs homozygous reference (‘AA’) mice at the top eQTL SNP for *Azi2* expression (Parker et al 2016; rs234453358) in the naïve CFW mice. **a** & **b** In the *Azi2* KO line, differential expression is performed on 16 striatum samples (wildtype = 5, heterozygote = 6, mutant = 5) and 15 VTA samples (wildtype = 3, heterozygote = 7, mutant = 5). **c** & **d** In the CFW mice, differential expression is performed on 17 striatum samples (‘AA’ = 5, ‘AG’ = 6, ‘GG’ = 6) and 17 VTA samples (‘AA’ = 6, ‘AG’ = 6, ‘GG’ = 5). Genes with FDR p-value < 0.05 are in shown in orange, and genes with FDR p-value < 0.1 are shown in black.

We sought to identify genome-wide changes in gene expression observed in the CFW mice after stratifying them by rs234453358, which was the peak eQTL SNP for *Azi2*. When comparing ‘AA’ to the ‘GG’ homozygotes, we identified five differentially expressed genomic features in the striatum at FDR < 0.1 (Figure 4c; Table S6). In the VTA, the same comparison identified only *Azi2* as being differentially expressed (Figure 4d; Table S6). When comparing the ‘AG’ to the ‘GG’ mice, we found five differentially expressed genomic features in the striatum (Figure S8c; Table S6) but none in the VTA (Figure S8d; Table S6). As discussed in the prior section, *Slc6a3* was not differentially expressed in any of these comparisons.

Finally, we performed PANTHER gene list analysis to reveal pathways implicated by these results (34). The mutant vs wildtype comparison in the striatum in the *Azi2* KO mice and the ‘GG’ vs ‘AA’ comparison in the striatum in the CFW mice both identified the Wnt signaling pathway (Table S6). Additional pathways that were identified include angiogenesis, Alzheimer disease-presenilin pathway, TGF-beta signaling pathway, metabotropic glutamate receptor group III pathway, cadherin signaling pathway, Notch signaling pathway, Huntington disease, nicotinic acetylcholine receptor signaling pathway, inflammation mediated by chemokine and cytokine signaling pathway, and cytoskeletal regulation by Rho GTPase (Table S6). A few genes had larger than five log_2_ fold changes but did not pass the FDR < 0.1 threshold, and thus were not considered as differentially expressed genes. Nevertheless, PANTHER gene list analysis showed that these genes are involved in pathways similar to those of differentially expressed genes (Table S7).

## Discussion

This purpose of the current study was to follow up on findings from the mouse GWAS by Parker et al (2016), which identified a locus that influenced the locomotor simulant response to methamphetamine and identified a co-localized *cis*-eQTL for the gene *Azi2*, which was hypothesized to be the casual variant. To experimentally test this hypothesis, we created a mutant allele of *Azi2*. The mutant allele reduced *Azi2* expression in the striatum (Figure 1), which was the tissue that showed an eQTL in Parker et al (2016) (20). Furthermore, most remaining transcripts were expected to be frameshifted and thus non-functional. Importantly, we observed significantly greater methamphetamine sensitivity in mutant *Azi2* mice (Figure 2F), supporting a role for this gene in the responses to methamphetamine. However, based on data from Parker et al (2016), we had predicted a positive relationship between *Azi2* expression and the locomotor stimulant response to methamphetamine. Instead, the mutant mice showed that reduced *Azi2* expression was associated with increased methamphetamine response. After the publication of Parker et al (2016), Liu et al (Liu et al., 2018) reported that the 3’UTR of *Azi2* regulated the expression of the dopamine transporter, which is the target of methamphetamine, and suggested that findings from Parker et al might be mediated by this mechanism. However, we did not observe any correlation between *Azi2* and *Sla6a3* in either the mutant *Azi2* mice or the CFW mice that harbor an eQTL for *Azi2* (Figure 3). Those observations do not support the hypothesis that *Azi2’s* effects are mediated by regulation of *Slc6a3* expression. While we did observe effects of the mutant *Azi2* allele on the expression of other genes (Figure 4), future studies will be needed to define the molecular pathway by which *Azi2* regulates sensitivity to methamphetamine.

A major conclusion from our work is that *Azi2* alters the locomotor response to methamphetamine. Although we did not exhaustively characterize these mice, they did not present any overt physical or behavioral abnormalities, and they did not show locomotor differences in the absence of methamphetamine administration (Figure 2d&e), suggesting that the effect of the mutation is at least somewhat specific to methamphetamine sensitivity. Parker et al (2016) found that the eQTL allele associated with greater *Azi2* expression was also associated with greater locomotor response to methamphetamine, whereas the current study found the opposite, namely that loss of *Azi2* was associated with greater response to methamphetamine. One possible explanation for this finding is that the effect of *Azi2* is modified by genetic background – the eQTL was observed in CFW mice, whereas the mutant allele was characterized using a C57BL/6J background. Consistent with this, we have previously reported that genetic background can induce directionally opposite effects of other mutant alleles (35). Another possibility is that the total loss of *Azi2* in the mutant line could have different consequences than the differential expression observed in the CFW mice. While both explanations are plausible, the difference in direction complicates our interpretation of the behavioral results and calls into question whether they should be considered to “replicate” or “recapitulate” the findings from Parker et al (2016).

One of the rationales for using an inbred C57BL/6J background was that expressing a mutant allele on an isogenic background would enhance our ability to detect an effect of the mutation, since it would remove other genetic differences that could be confounding. It is notable that despite this advantage, a relatively large sample size was required to obtain significant results. We observed a similar result in a prior study in which we examined a mutant allele of the gene *Csmd1* (19), which had been implicated by a separate mouse GWAS (36). While these two examples do not represent a large enough sample to draw general conclusions, it may be that genes identified using mouse GWAS can have relatively subtle effects that require sample sizes that are larger than those often employed when examining mutant mice. Because studies like these do not use the alleles identified in the GWAS, power analyses are difficult because the expected effect size is unknown. Our observations imply that future studies following up on mouse GWAS should consider using relatively large samples (in this case more than 100 total subjects) before drawing conclusions.

The goal of our study was to determine whether *Azi2* is the gene responsible for the association detected by Parker et al (2016). That locus contained a second candidate gene, COX assembly mitochondrial protein 1 (***Cmc1***), which could also have contained regulatory variants for other nearby genes. it is possible that the locus harbored multiple causal variants. Studies such as ours, even when perfectly successful, are not able to refute the possibility that multiple genes contribute to a given association.

In an effort to define the causal pathway by which *Azi2* might alter sensitivity to the locomotor effects of methamphetamine, we tested the hypothesis that the 3’UTR of *Azi2* regulates *Slc6a3*, as described by Liu et al (Liu et al., 2018). Using a luciferase reporter assay in the human neuroblastoma SK-N-AS cells, Liu et al (2018) identified *Azi2* 3’UTR as a putative downregulator of the promoter activity in only one allele of a dinucleotide polymorphism in Intron 1 of *SLC6A3*, but not the other (Liu et al., 2018). It is possible that this allele specificity is the reason for the lack of correlation we observed between the 3’UTR of *Azi2* and the expression of *Slc6a3*. However, Liu et al (2018) did not detect any *SLC6A3* allele-dependence in the downregulation of endogenous *SLC6A3* mRNA level by *AZI2* 3’UTR in the human neuroblastoma BE(2)-M17 cells. Furthermore, Northern blot and qRT-PCR results on *Azi2*/ *Azi2* 3’UTR and *Slc6a3* expression in the VTA of alcohol preferring and non-preferring rats did not distinguish the two alleles of *Slc6a3* (Liu et al., 2018). Thus, we investigated the correlation between *Azi2*/ *Azi2* 3’UTR and *Slc6a3* expression in both our *Azi2* KO mice and naïve CFW mice, intending to replicate the experiments performed on alcohol preferring and non-preferring rats (Liu et al., 2018). Our results do not contradict the findings of Liu et al (2018) but they strongly suggest that *Azi2*’s actions observed in our studies are not mediated by *Slc6a3*.

Although we did not find evidence to support a role for *Slc6a3* in the effects of *Azi2*, we did identify a number of other genes that were differentially expressed in both the *Azi2* KO line and in the CFW mice (Figure 4). One differentially expressed gene, *Slc16a6*, is mapped to the metabotropic glutamate receptor group III pathway. Another gene, *Myh1*, is mapped to the nicotinic acetylcholine receptor signaling pathway. Future studies should investigate whether these genes and their involvement in synaptic signaling pathways might hold clues about the relationship between *Azi2* and sensitivity to methamphetamine.

Our study is not without limitations. For example, the creation of the mutant allele using CRISPR/Cas9 could have induced unintended off-target mutations. We backcrossed the mutant allele for several generations, and unless putative off-target mutations were nearby and thus linked to the mutant allele, they should have segregated independently, since all experiments uses wildtype littermate controls. Nevertheless, it is technically possible that a linked, off-target mutation may have interfered with our results. We also did not characterize the effect of the *Azi2* mutation on other doses of methamphetamine nor did we examine other behavioral traits of these mice, which might have provided clues about the possible role of *Azi2* in substance abuse-related traits. Finally, we conducted all of the studies intended to examine the role of *Slc6a3* in naïve C57BL/6J and CFW mice, whereas Liu et al (2018) examined human cell lines, human postmortem dopamine neurons, and alcohol preferring and alcohol non-preferring rats (Liu et al., 2018). Thus, our conclusions about the lack of correlation between *Azi2* and *Slc6a3* only apply to the mouse systems that we examined.

The present study is notable because it remains rare to experimentally test specific genes identified using model organism or human GWAS. Our results highlight the potential for such studies, including their ability to contribute to a molecular understanding of how a specific gene influences a specific trait, which is essential for deriving new biological insights from GWAS results. However, our results also illustrate challenges, including the choice of background, and the criteria needed to claim replication.

## Author Contributions

AAP and XZ designed the study, oversaw data collection and analysis, and co-wrote the manuscript. XZ designed the CRISPR/Cas9 knockout scheme, maintained mouse colonies, conducted behavioral experiments, collected brain tissues, extracted RNA and DNA from mice, genotyped the mice, performed qRT-PCR, and analyzed the behavioral and gene expression data under supervision of AAP. ABL provided help and training with mouse handling, mouse behavioral tests and brain dissection. PM performed quality control of RNA and prepared cDNA libraries for RNA-Seq under supervision of FT.

