## Supplementary figures and images for "Functional validation of a finding from a mouse genome-wide association study demonstrates that a mutant allele of *Azi2* alters sensitivity to methamphetamine"

### Figure S1

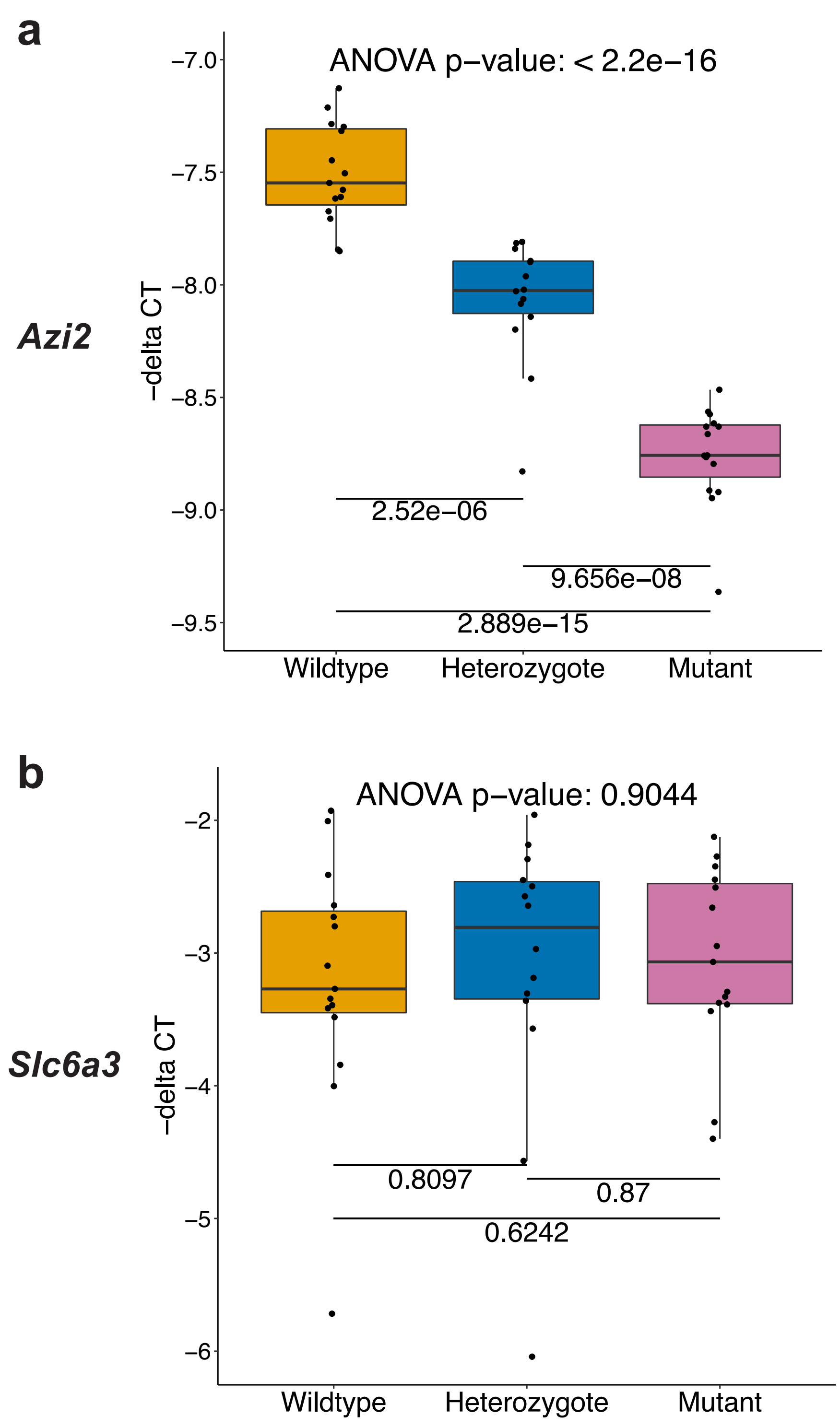

### Figure S2

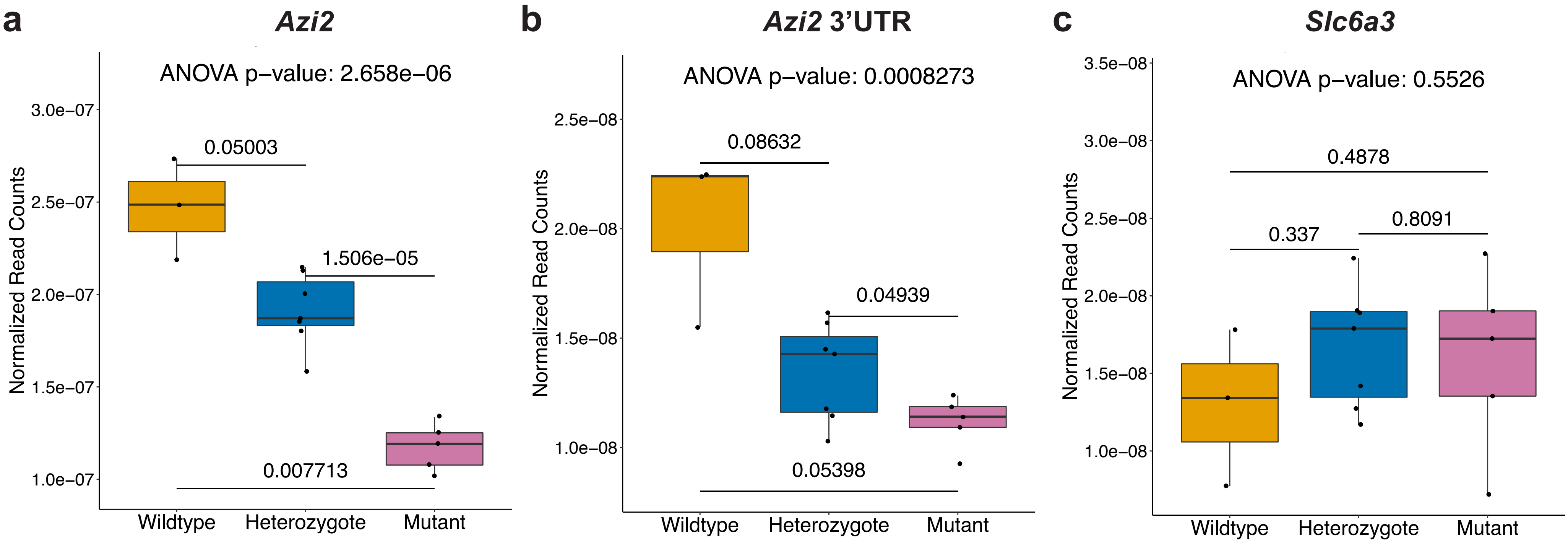

### Figure S3

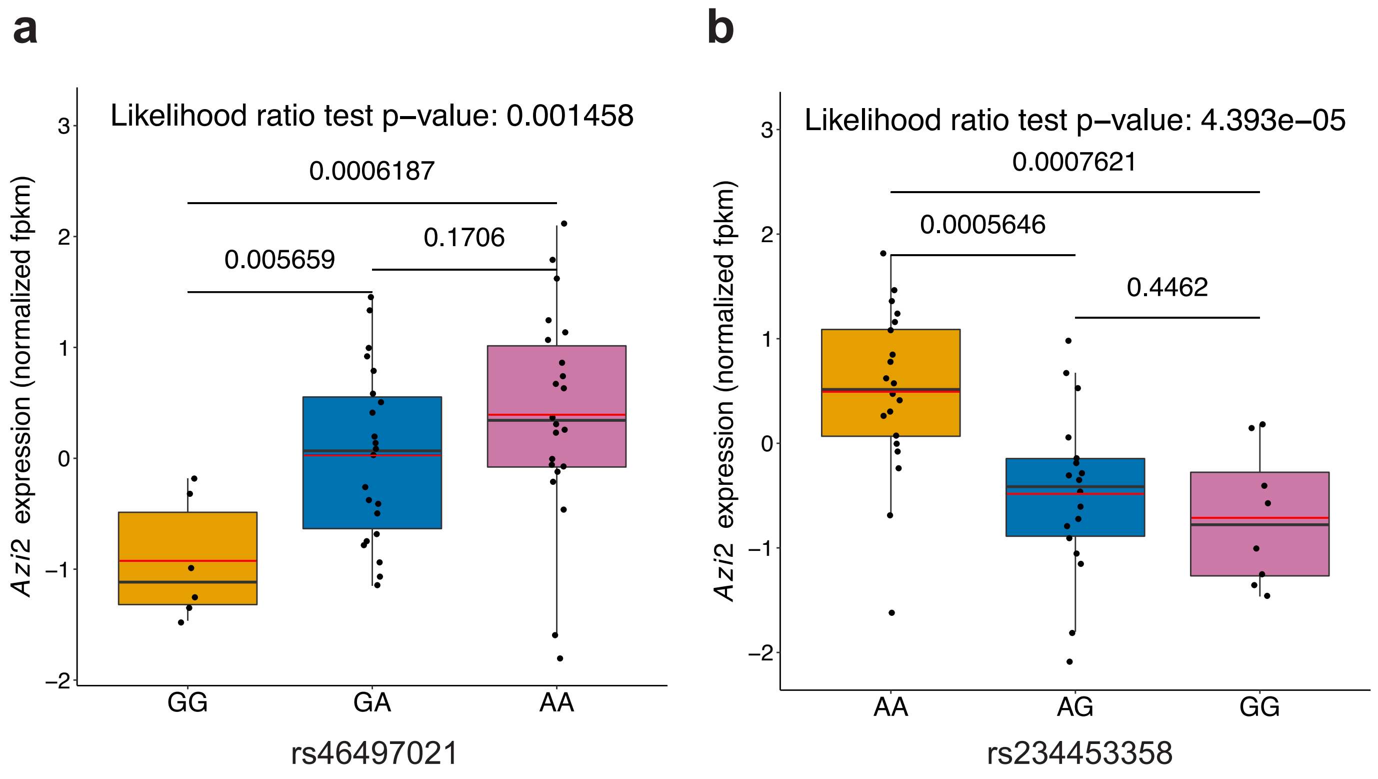

### Figure S4

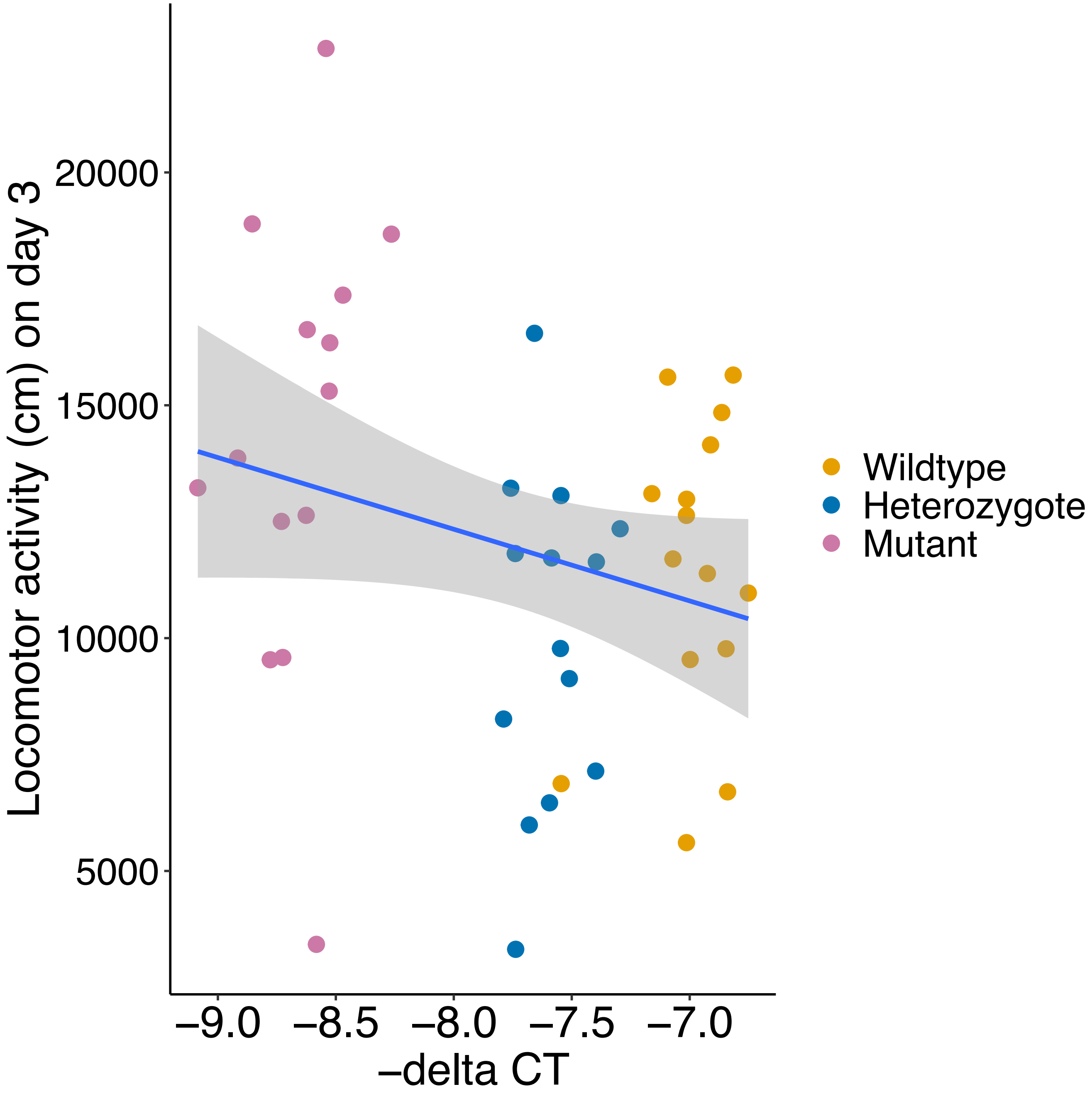

### Figure S5

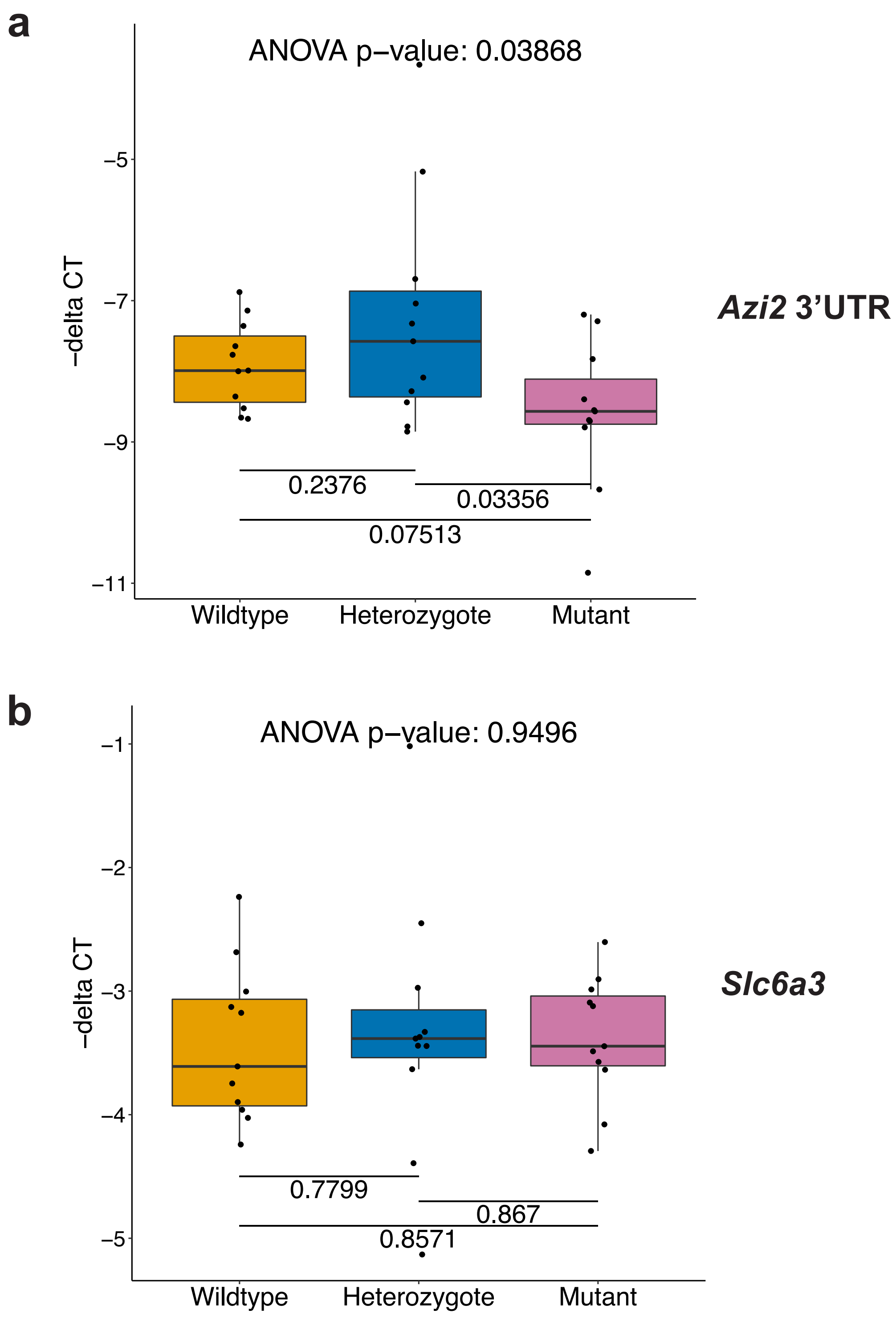

### Figure S6

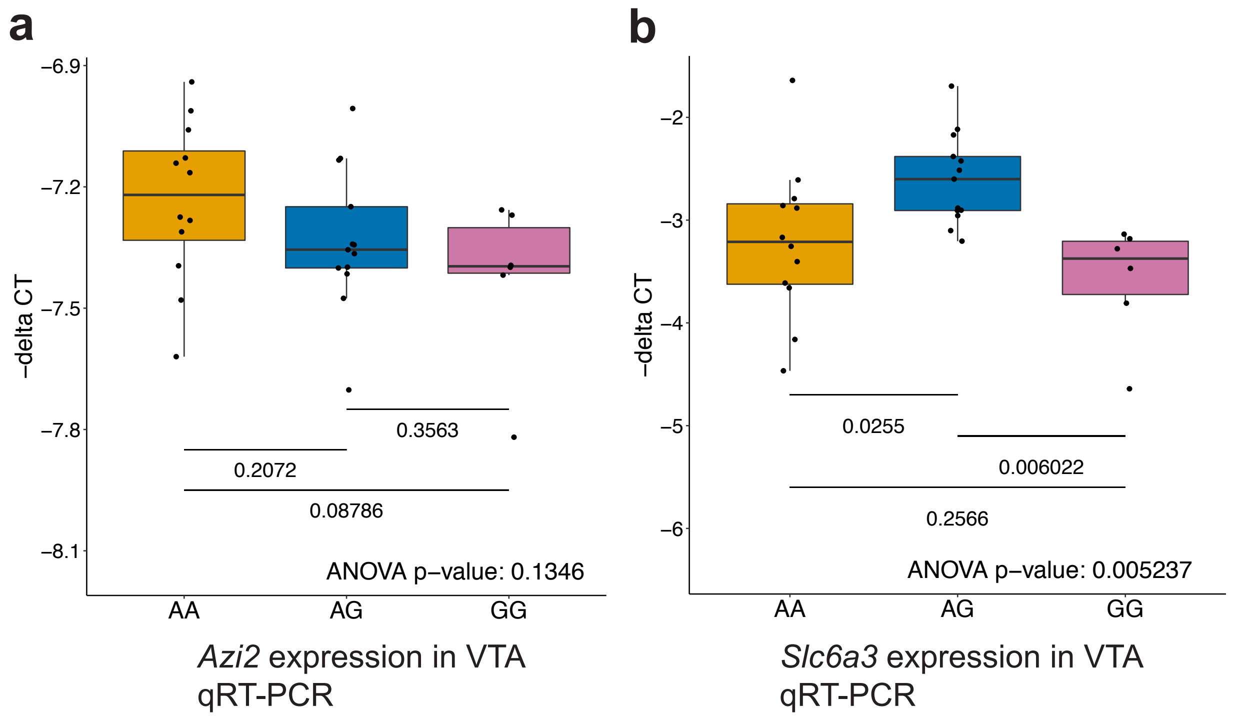

### Figure S7

**c**

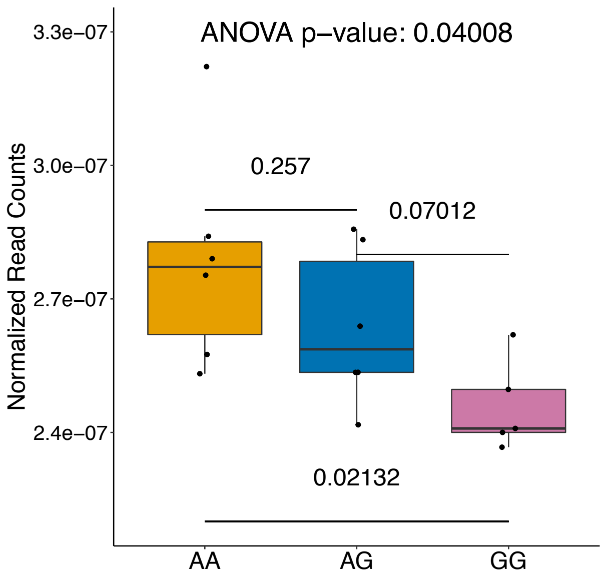


*Azi2* expression in VTA RNA-Seq

**d**

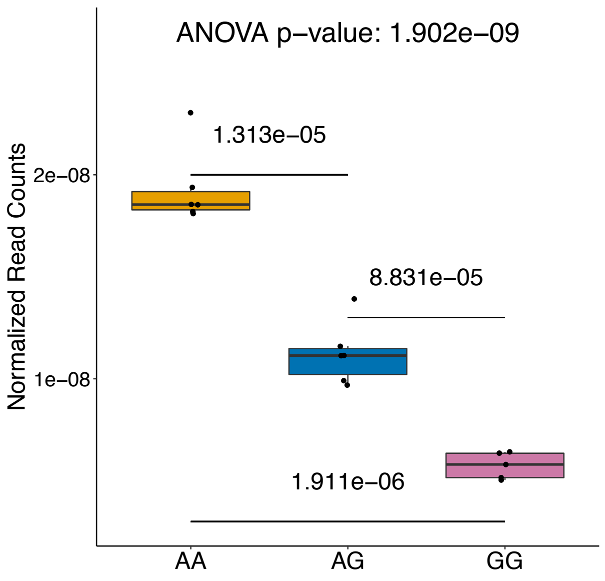


*Azi2* 3'UTR expression in VTA RNA-Seq

**e**

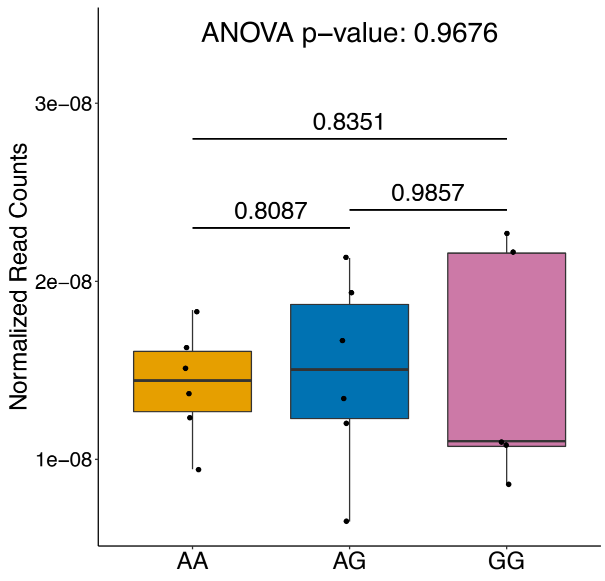


*Slc6a3* expression in VTA RNA-Seq

### Figure S8

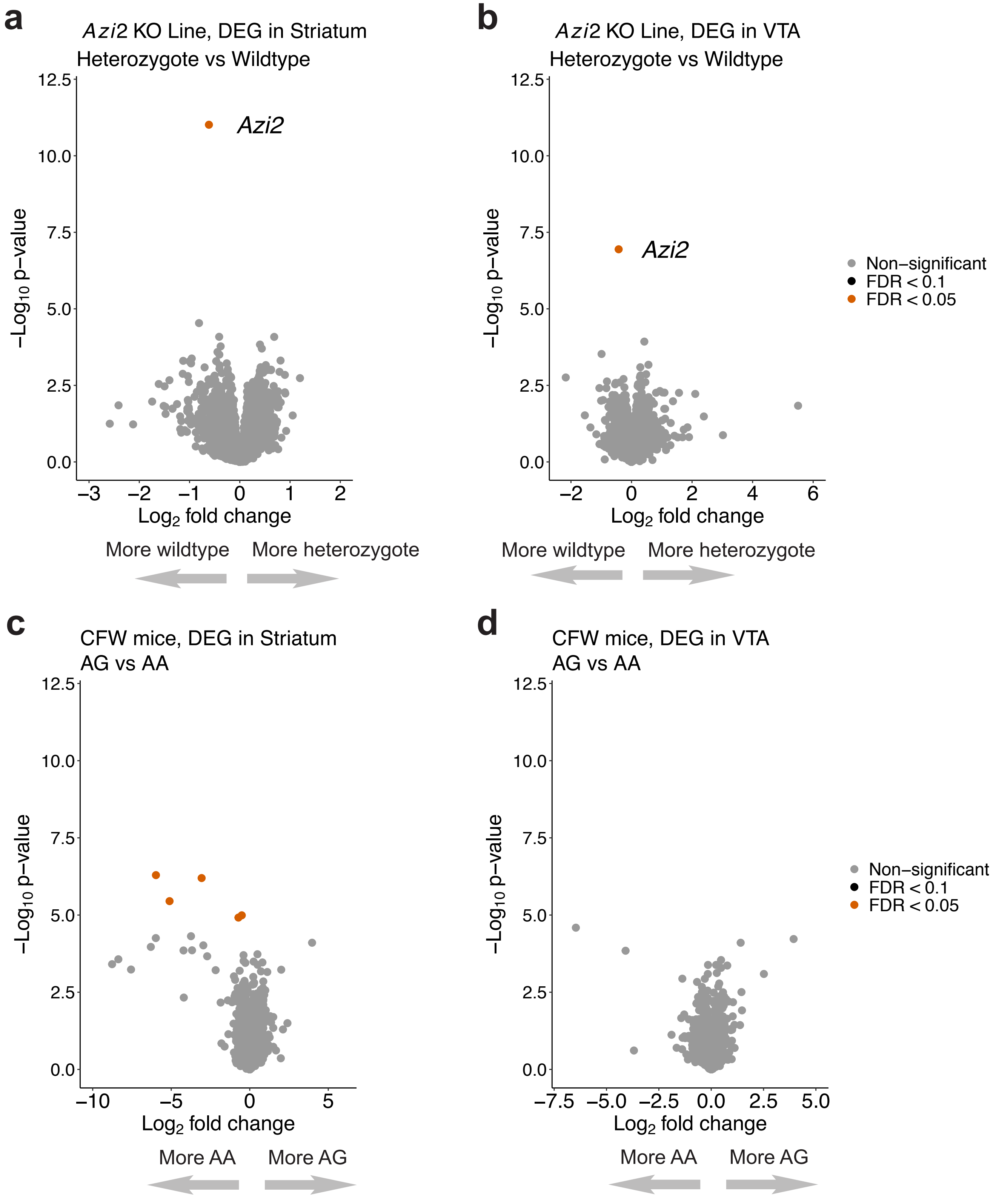
