## Supplementary figure legends for "Functional validation of a finding from a mouse genome-wide association study demonstrates that a mutant allele of *Azi2* alters sensitivity to methamphetamine"

**Figure S1a&b. qRT-PCR data did not support the hypothesis that *Azi2* downregulates *Slc6a3* in the VTA in the *Azi2* KO mice.** We used 44 *Azi2* KO mice (wildtype = 15, heterozygote = 14, mutant = 15) for examining *Azi2* and *Slc6a3* expression, measured by qRT-PCR. **a** Mutant *Azi2* KO mice showed a lower *Azi2* expression level in the VTA (*F*(2,41) = 102.68, p < 2.2×10^-16^). All between-group comparisons were significant (wildtype vs heterozygote *t*(25.063) = -6.0496, p = 2.52×10^-6^; heterozygote vs. mutant *t*(24.898) = -7.4014, p = 9.656×10^-8^; wildtype vs mutant *t*(27.995) = -15.497, p = 2.889×10^-15^). **b** Genotype did not affect *Slc6a3* expression (*F*(2,41) = 0.1007, p = 0.9044). All between-group comparisons were not significant (wildtype vs. heterozygote *t*(25.676) = 0.24334, p = 0.8097; heterozygote vs mutant *t*(21.761) = 0.16561, p = 0.87; wildtype vs mutant *t*(25.853) = 0.49582, p = 0.6242).

**Figure S2a-c. RNA-Seq data did not support the hypothesis that *Azi2*/*Azi2* 3’UTR down-regulates *Slc6a3* in the VTA in the *Azi2* KO mice.** Fifteen *Azi2* KO mice (wildtype = 3, heterozygote= 7, mutant= 5) for examining the correlation between *Azi2*/*Azi2* 3’UTR and *Slc6a3* normalized read counts measured by RNA-Seq. **a** & **b** Mutant *Azi2* KO mice showed a lower normalized read count mapped to *Azi2* in the VTA (*F*(2,12) = 44.978, p = 2.658×10^-6^) and to *Azi2* 3’UTR (*F*(2,12) = 13.583, p = 0.0008273). **a** Wildtype vs heterozygote comparison was trending significant (*t*(2.9714) = 3.1991, p = 0.05003); the other genotype comparisons were highly significant (heterozygote vs mutant *t*(9.965) = 7.7963, p = 1.506×10^-5^; wildtype vs mutant *t*(2.5547) = 7.749, p = 0.007713). **b** All between-group comparisons have p < 0.1 (wildtype vs heterozygote *t*(2.5699) = 2.7145, p = 0.08632; heterozygote vs mutant *t*(9.4254) = 2.2541, p = 0.04939; wildtype vs mutant *t*(2.2148) = 3.7787, p = 0.05398). **c** Deletion did not affect *Slc6a3* expression (*F*(2,12) = 0.6234, p = 0.5526). All between-group comparisons were not significant (wildtype vs heterozygote *t*(3.0881) = -1.1342, p = 0.337; heterozygote vs mutant *t*(6.4731) = 0.25173, p = 0.8091; wildtype vs mutant *t*(4.9537) = -0.74919, p = 0.4878).

**Figure S3a&b. *Azi2* expression in the striatum of CFW mice published in Parker et al (2016). a & b** *Azi2* expression at the GWAS top SNP (rs46497021) and at the eQTL top SNP (rs234453358) for locomotor activity on day three when each mouse received 1.5 mg/kg methamphetamine injection. Dosages < 0.2 are coded homozygous reference, dosages > 0.8 and <1.2 heterozygous, dosages > 1.8 homozygous alternative in panels **a** & **b**. Likelihood ratio tests of nested models were performed on all included dosages. **a** Mice with the homozygous alternative AA genotype at the GWAS top SNP have significantly higher *Azi2* expression in the striatum (‘GG’ n = 6, ‘GA’ n = 22, ‘AA’ n = 22; likelihood ratio test nested models, *X*^2^(1,2) = 10.131, p = 0.001458). ‘GG’ vs ‘GA’ (*t*(11.266) = -3.4078, p = 0.005659) and ‘GG’ vs ‘AA’ (*t*(14.541) = -4.3445, p = 0.0006187) comparisons were significant. ‘GA’ vs ‘AA’ comparison was not significant (*t*(40.279) = -1.3952, p = 0.1706). **b** Mice with the homozygous alternative GG genotype at the eQTL top SNP have significantly higher *Azi2* expression in the striatum (‘AA’ n = 20, ‘AG’ n = 18, ‘GG’ n = 8; likelihood ratio test nested models, *X*^2^(1,2) = 16.694, p = 4.393×10^-05^). ‘AA’ vs ‘AG’ (*t*(35.657) = 3.7867, p = 0.0005646) and ‘AA’ vs ‘GG’ (*t*(15.876) = 4.1517, p-value = 0.0007621) comparisons were significant. ‘AG’ vs ‘GG’ comparison was not significant (*t*(16.325) = 0.78064, p = 0.4462).

**Figure S4. Higher *Azi2* expression in the striatum corresponds to lower locomotor activity on day 3 in *Azi2* KO mice (*r*(41) = -0.26482, p = 0.08613).** We plotted a subset of the *Azi2* KO mice that were used in the locomotor activity test that were also measured in qRT-PCT (n=43).

**Figure S5a&b. qRT-PCR data did not support the hypothesis that *Azi2*/*Azi2* 3’UTR down-regulates *Slc6a3* in the VTA in the *Azi2* KO mice.** We used 33 *Azi2* KO mice (wildtype = 11, heterozygote = 11, mutant = 11) for examining *Azi2* 3’UTR and *Slc6a3* expression, measured by qRT-PCR. **a** Genotype effect on *Azi2* 3’UTR expression was only moderately significant in the VTA (*F*(2,30) = 3.6319, p = 0.03868). Wildtype vs heterozygote comparison (*t*(12.875) = 1.2385, p = 0.2376) and wildtype vs mutant comparison (*t*(16.334) = -1.9008, p = 0.07513) were not significant. Heterozygote vs mutant comparison was significant (*t*(17.035) = -2.3118, p = 0.03356). **b** Genotype did not affect *Slc6a3* expression (*F*(2,30) = 0.0518, p = 0.9496). All between-group comparisons were not significant (wildtype vs. heterozygote *t*(16.49) = 0.28401, p = 0.7799; heterozygote vs mutant *t*(14.535) = -0.17051, p = 0.867; wildtype vs mutant *t*(19.142) = 0.18256, p = 0.8571).

**Figure S6a&b. qRT-PCR data showed that *Azi2*/*Azi2* 3’UTR did not down-regulate *Slc6a3* regulation in the VTA in the CFW naïve mice. a** *Azi2* expression in the VTA, measured by qRT-PCR, did not show a significant genotype effect at the eQTL top SNP for *Azi2* expression (rs234453358; *F*(2,28) = 2.1559, p = 0.1346; ‘AA’ = 12, ‘AG’ = 13, ‘GG’ = 6). All between-group comparisons were not significant (‘AA’ vs ‘AG’ *t*(22.034) = -1.2995, p = 0.2072; ‘AG’ vs ‘GG’ *t*(8.5552) = -0.97499, p = 0.3563; ‘AA’ vs ‘GG’ *t*(9.8541) = = -1.8944, p = 0.08786). **b** Genotype was significant in *Slc6a3* expression in the VTA (*F*(2,28) = 6.3729, p = 0.005237); however, the genotype effect was not additive. AA’ vs ‘AG’ (*t*(17.563) = 2.4408, p = 0.0255) and ‘AG’ vs ‘GG’ (*t*(7.8689) = -3.7229, p = 0.006022) comparisons were significant, while ‘AA’ vs ‘GG’ comparison was not (*t*(12.881) = -1.1871, p = 0.2566).

**Figure S7a-c. RNA-Seq data showed that *Azi2*/*Azi2* 3’UTR did not down-regulate *Slc6a3* regulation in the VTA in the CFW naïve mice.** While *Azi2* (*F*(2,14) = 4.0836, p = 0.04008) and *Azi2* 3’UTR (*F*(2,14) = 116.29, p = 1.902×10^-9^) normalized read counts showed significant decrease at the homozygous ‘GG’ allele, *Slc6a3* normalized read counts exhibited no genotype effect at this SNP (*F*(2,14) = 0.033, p = 0.9676). **a** ‘AA’ vs ‘AG’ (*t*(9.065) = 1.2097, p = 0.257) and ‘AG’ vs ‘GG’ (*t*(8.1698) = 2.0824, p = 0.07012) comparisons were not significant. ‘AA’ vs ‘GG’ comparison was significant (*t*(6.9047) = 2.9642, p = 0.02132). **b** All between-group comparisons were significant (‘AA’ vs ‘AG’ *t*(9.5467) = 8.1646, p = 1.313×10^-5^; ‘AG’ vs ‘GG’ *t*(7.0148) = 8.0254, p-value = 8.831×10^-5^; ‘AA’ vs ‘GG’ *t*(6.3521) = 16.444, p = 1.911×10^-6^). **c** All between-group comparisons were not significant (‘AA’ vs ‘AG’ *t*(8.0372) = -0.25024, p = 0.8087; ‘AG’ vs ‘GG’ *t*(7.7373) = -0.018508, p = 0.9857; ‘AA’ vs ‘GG’ *t*(5.4762) = -0.21828, p = 0.8351).

**Figure S8a-d. *Azi2* (ENSMUSG00000039285.12) was differentially expressed between heterozygous vs wildtype mice in the *Azi2* KO line, but not between heterozygous alternative (‘AG’) vs homozygous reference (‘AA’) mice at the top eQTL SNP for *Azi2* expression (Parker et al 2016; rs234453358**) **in the naïve CFW mice.**  **a** & **b** In the *Azi2* KO line, differential expression is performed on 16 striatum samples (wildtype = 5, heterozygote = 6, mutant = 5) and 15 VTA samples (wildtype = 3, heterozygote = 7, mutant = 5).  **c** & **d** In the CFW mice, differential expression is performed on 17 striatum samples (‘AA’ = 5, ‘AG’ = 6, ‘GG’ = 6) and 17 VTA samples (‘AA’ = 6, ‘AG’ = 6, ‘GG’ = 5). Genes with FDR p-value < 0.05 are shown in orange, and genes with FDR p-value < 0.1 are shown in black.
